# Behavioral, neural and ultrastructural alterations in a graded-dose 6-OHDA mouse model of early-stage Parkinson’s disease

**DOI:** 10.1101/2023.03.08.531684

**Authors:** Andrea Slézia, Panna Hegedüs, Evgeniia Rusina, Katalin Lengyel, Nicola Solari, Attila Kaszas, Diána Balázsfi, Boris Botzanowski, Emma Acerbo, Florian Missey, Adam Williamson, Balázs Hangya

## Abstract

Studying animal models furthers our understanding of Parkinson’s disease (PD) pathophysiology by providing tools to investigate detailed molecular, cellular and circuit functions. Different versions of the neurotoxin-based 6-hydroxydopamine (6-OHDA) model of PD have been widely used in rats. However, these models typically assess the result of extensive and definitive dopaminergic lesions that reflect a late stage of PD, leading to a paucity of studies and a consequential gap of knowledge regarding initial stages, in which early interventions would be possible. Additionally, the better availability of genetic tools increasingly shifts the focus of research from rats to mice, but few mouse PD models are available yet. To address these, we characterize here the behavioral, neuronal and ultrastructural features of a graded-dose unilateral, single-injection, striatal 6-OHDA model in mice, focusing on early-stage changes within the first two weeks of lesion induction. We observed early onset, dose-dependent impairments of overall locomotion without substantial deterioration of motor coordination. In accordance, histological evaluation demonstrated a partial, dose-dependent loss of dopaminergic neurons of substantia nigra pars compacta (SNc). Furthermore, electron microscopic analysis revealed degenerative ultrastructural changes in SNc dopaminergic neurons. Our results show that mild ultrastructural and cellular degradation of dopaminergic neurons of the SNc can lead to certain motor deficits shortly after unilateral striatal lesions, suggesting that a unilateral dose-dependent intrastriatal 6-OHDA lesion protocol can serve as a successful model of the early stages of Parkinson’s disease in mice.

**Highlights:**

- Unilateral striatal 6-OHDA injection caused an early onset dose-dependent behavioral deficit in mice
- Behavioral deficit manifested as mild impairment of locomotion and locomotive movement initialization
- Behavioral changes were accompanied by moderate, dose-dependent loss of SNc dopaminergic neurons
- Behavioral changes were paralleled by ultrastructural alterations of SNc dopaminergic neurons

## 1. Introduction

Parkinson’s disease (PD) is the second most common neurodegenerative disorder affecting more than seven million people worldwide(*1–4*). PD is a progressive neurodegenerative disorder; it is dominated by motor symptoms, while non-motor problems including autonomic dysfunctions and cognitive impairment are not uncommon among PD patients(*5*). The motor deficits include resting tremor, bradykinesia, rigidity, and postural instability, caused by the selective, large scale, irreversible degeneration of dopaminergic (DAergic) neurons mostly located in the substantia nigra pars compacta (SNc) of the midbrain. This gradually progressing, selective SNc DAergic cell loss leads to a progressive reduction of dopamine (DA) concentration in the striatum(*6, 7*). To date, there are no disease-modifying therapies that slow, halt, or reverse the progression of PD(*8*).

Importantly, the above-mentioned symptoms become apparent, and thus the diagnosis is usually made, at a rather advanced stage of neurodegeneration; on average, 50-80% of nigral DA neurons are already lost by that time(*6, 9, 10*). Accordingly, most of the classic literature based on animal experiments – monkey and rodent models – and human studies reflect late, chronic, dopamine-depleted states when compensatory mechanisms are already in place in most related neural network functions(*2, 11, 12*). In the past decades, however, early stages of the disease have received increasing focus(*13–15*). Neurodegenerative processes start to play a role in the manifestation of the disease decades before the diagnosis(*3, 16*), including disrupted synaptic and cellular plasticity and consequential impairments of functional connectivity that eventually leads to altered motor system dynamics associated with PD. Detailed behavioral analysis can help to better understand these network changes and aid PD diagnosis at an earlier timepoint, possibly allowing a timely clinical intervention(*15, 17, 18*).

Therefore, it is important to develop research models that can help us pinpoint mechanisms of early changes in brain function during disease. In particular, graded models will allow a more differentiated picture of disease symptoms by enabling us to study both mild and severe lesions and correlated pathophysiological features at different stages of progression (*19, 20*), potentially opening the way to novel neuroprotective therapies and early intervention or prevention of PD(*21*). In addition, it could help the development of novel diagnostic methods that, in turn, could lead to preventing or stopping disease manifestation in human patients(*22*). However, animal studies on early neuropathological and behavioral changes are sparse, and little information is available on simultaneous behavioral, histological, and cellular ultrastructural changes at the early stages of PD, limiting our mechanistic understanding of early-stage disfunctions(*23–28*).

Here, we studied the histological and behavioral alterations that characterize the early stages of PD in a graded 6-OHDA mouse model. Our results from open field and rotarod behavioral assays show that mild motor impairments are detectable already at very early stages, even one week after a single low dose of striatal 6-OHDA injection, typically not explored in 6-OHDA models of PD. However, motor impairments mostly manifested in general horizontal locomotion and initiation of explorative locomotion, rather than in motor coordination, which was regularly found affected by PD at later stages(*12, 29*). While the mild impairment in exploratory motor behavior was maintained and aggravated after the second week in a dose-dependent manner, motor coordination tested in the rotarod remained similar across 6-OHDA-injected and sham-operated mice. Motor symptoms were accompanied by a significant dose-dependent loss of DAergic cells in the SNc already after one week post injection. Additionally, ultrastructural changes in SNc DAergic neurons, such as the degeneration of mitochondria (swelling, structural changes of the lamellae) and the endoplasmic reticulum were detected by electron microscopy.

These findings demonstrate the presence of early-onset structural and behavioral impairments in a single-dose, unilateral striatal 6-OHDA mouse model of PD, suggesting that the partial lesion protocol we developed might be suitable for investigating mechanistic neuropathological changes in early phases of the disease, holding the promise for developing early diagnostics and interventions saving many disability-adjusted life years, and potentially even paving the way towards disease-modifying therapies.

## 2. Materials and methods

### 2.1 Animals

Sixty adult male C57BL6/J mice were used for the experiments. Mice were kept on a 12 h light/dark cycle with food and water available *ad libitum*. All experiments were approved by the Committee for Scientific Ethics of Animal Research of the National Food Chain Safety Office (PE/EA/784-7/2019) and were performed according to the guidelines of the institutional ethical code and the Hungarian Act of Animal Care and Experimentation (1998; XXVIII, section 243/1998, renewed in 40/2013) in accordance with the European Directive 86/609/CEE and modified according to the Directive 2010/63/EU.

### 2.2 Surgical procedure

Animals were briefly sedated with isoflurane (Forane-abbvie Hungaropharma, Hungary) in a sealed container (15-30 sec) to reduce the stress associated with anesthesia. Buprenorphine analgesic was administered subcutaneously (Bupaq, 10%, 0.05 - 0.2 mg / kg, Primavet, Austria). Surgery was performed under ketamine-xylazine anesthesia (i.p. ketamine: 50 mg / kg; xylazine: 20 mg / kg, Medicus Partner, Netherlands). The depth of the anesthesia was monitored by checking reflexes, breathing frequency, and whisker movements. When adequate anesthesia was reached, animals were placed into a stereotaxic apparatus (Stoelting, UK).

The skin over the calvaria was shaved and disinfected with Betadine (EGIS, Hungary). Eyes were protected from dehydration and strong light with rodent eye ointment (Bausch & Lomb, Germany), repeated during surgery as necessary. The skin and connective tissues over the skull were infused by local anesthetics (Ropivacaine s.c., Braun Melsungen AG., Germany), and a midline scalp incision was made. Subcutaneous tissues were removed, the skull was cleaned, and the dura mater was removed. The animal’s head was adjusted so that Bregma and Lambda were in the same horizontal plane. A craniotomy was made over the dorsal striatum (stereotaxic coordinates from bregma: anteroposterior 0.6 mm and mediolateral 1.8 mm). A borosilicate capillary pipette (Sutter Instrument, Germany) was lowered to the target area (dorsoventral −2.0 mm from the cortical surface) and 6-OHDA (Sigma-Aldrich, Hungary) was injected by a syringe pump (World Precision Instruments, UK), delivering 1μl of 6-OHDA solution diluted in 0.02% ascorbic acid (Sigma-Aldrich, Hungary) with a speed of 1μl/min in either a low, medium or high dose (2.5μg/μl, 5μg/μl or 8μg/μl 6-OHDA, respectively; Fig.1A-B)(*19*). For sham surgeries, we used the same stereotaxic coordinates and injected 1μl of 0.02% ascorbic acid. The pipette was removed after 3-4 minutes waiting time and the wound edges were stitched or closed by Vetbond surgical glue (3M, Hungary). The animals were continuously monitored and observed for the signs of pain, distress, or neurological complications for a one-week recovery time following the surgery.

**Figure 1.**
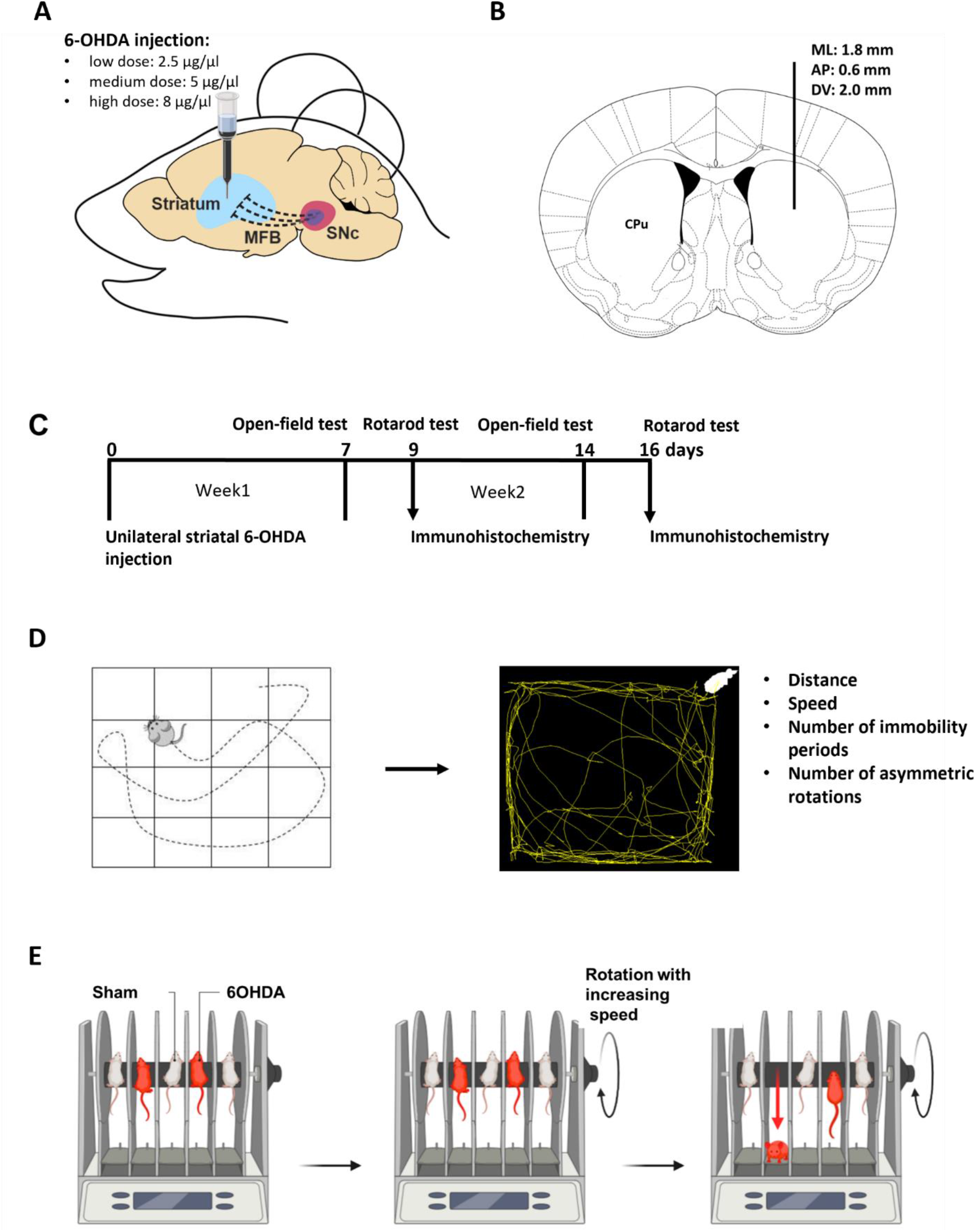
Methodological overview. A, Schematic of the unilateral intrastriatal 6-OHDA injection. B, Intrastriatal injection site marked in the mouse brain atlas in the coronal plane. C, Timeline of the experiment. Open field test was performed 7 and 14 days after 6-OHDA injection. Rotarod test was performed 9 and 16 days after the lesion. Half of the treated animals per group (LD, n = 5; MD, n = 4; HD, n = 4) were sacrificed and processed for immunohistochemistry after the first rotarod test, while the remaining mice (LD, n = 6; MD, n = 5; HD, n = 5) underwent immunohistochemistry procedures after an additional week of testing. D, Schematic of open-field test protocol and analysis. E, Schematic for rotarod test protocol.

Standard surgical procedures were carefully used to minimize animal suffering, brain tissue damage, and risk of infection. Operated mice were kept in separate cages. They were checked regularly in a 48-hour-postoperative period. Bodyweight change was monitored for three days after the surgery.

### 2.3 Evaluation of locomotor activity, exploration, motor coordination, and voluntary movement

#### 2.3.1 Open field test – Measurement of spontaneous locomotor activity, movement initiation, and exploration

Mice were placed separately in a 40 x 40 x 30 cm plastic box with a 4 x 4 square mesh on its bottom and were video-recorded for 5 minutes (Fig.1C-D). The following parameters of locomotion were extracted: (1) distance in cm, (2) speed in cm/min, (3) number of immobility periods, and (4) number of asymmetric rotations. The movement trajectories of mice were detected in the videos and were further analyzed by Image J software and Animal Tracker plugin(*30*), where overall distance covered, speed, the number of line crossings and spontaneous rotations were measured. Immobility periods were defined as events when mice were in the same cm^2^ for at least 2s.

#### 2.3.2 Rotarod test – Testing motor coordination

Motor coordination was tested with rotarod equipment (Bioseb, France; Fig.1E). To evaluate motor coordination and balance, the animals were placed on an accelerating rotating cylinder and the latency until falling from the rotating rod was measured. We used an accelerating speed from 4 RPM to 40 RPM. The tests lasted for a maximum of 3 minutes and were repeated three times for each mouse with a five-minute break after each session. During breaks, mice were placed back in their own cages.

#### 2.3.3 Statistical analysis of behavioral tests

Statistical analyses were performed using one-way standard analysis of variance by using the GraphPad Prism 6.01 software (GraphPad Inc., San Diego, CA, USA). When a significant difference was found between groups, we performed Tukey’s honest significant difference (HSD) post hoc multiple comparison test to identify pairwise differences. For all tests, p< 0.05 was considered significant.

### 2.4 Evaluation of neurodegenerative processes

54 male, adult, C57BL/6 mice were used in the experiments. Histological evaluations were made in all animals that participated in the behavioral tests. After the first week of behavioral experiments, n = 11 low-dose, n = 9 medium dose, n = 9 high-dose, and n = 6 sham animals were analyzed. Following the second week, n = 6 low-dose, n = 5 medium dose, n = 5 high-dose, and n = 6 sham animals were examined. All sections of all 54 animals were examined by light microscopy. Cell counting was performed in 2 animals from each group.

The regions of interest were selected using the mouse brain atlas(*31–33*) for further analysis of labeled TH+ neurons by immunohistochemistry and electron microscopy. For electron microscopic imaging, SNc and striatum were localized in 50 µm coronal sections by using the mouse brain atlas as a reference. SNc and striatal tissue samples were dissected with a surgical scalpel from each section. Tissue samples were then mounted on epoxy blocks for ultramicrotome sectioning.(*31–33*)

#### 2.4.1 Immunohistochemistry – visualizing DAergic cell loss

Mice were euthanized with an overdose of pentobarbital (100 mg/kg i.p.). Animals were transcardially perfused first with saline, then with 150 ml of a fixative solution containing 4% paraformaldehyde (PFA) in 0.1 M phosphate buffer (PB). Tissue blocks were cut on a Vibratome (Leica VT1200S, Leica Microsystems, Wetzlar, Germany) into 50 µm coronal sections.

After slicing and extensive washing in 0.1 M PB (3 times for 10 min), the 50-μm-thick sections were incubated in 30% sucrose overnight, freeze-thawing over liquid nitrogen four times, then processed for immunohistology. All following washing steps and dilutions of the antibodies were done in 0.05 M TBS buffer (pH = 7.4). After extensive washing in TBS (3 times for 10 min), the sections were blocked in 5% normal goat serum for 45 min and then incubated in the primary antibody for a minimum of 48 h at 4°C (mouse anti-tyrosine hydroxylase monoclonal antibody recognizing an epitope in the mid-portion of the rat TH protein; dilution: 1:8000; Product No: 22941; ImmunoStar). After incubating with the primary antibody, the sections were treated with biotinylated donkey-anti-mouse IgG (1:300; Vector Laboratories) for 2 h. Next, the sections were incubated with avidin biotinylated–horseradish peroxidase complex (1:500; Elite ABC; Vector Laboratories) for 1.5 h. The immunoperoxidase reactions were finally developed using 3,3′-diaminobenzidine 4HCl (DAB) as the chromogen. Sections were dehydrated in xylol, then mounted in chrome-gelatin and covered by DePeX. (*34*).

#### 2.4.2 Neuronal counts – quantifying DAergic cell loss

For quantitative analysis, we selected sections in which corresponding structures appeared symmetrically across the hemispheres at each AP level to avoid potential biases due to asymmetric sectioning. Specifically, the symmetry of dorsal and ventral hippocampi, nuclei of thalami (thalamic reticular nucleus, laterodorsal thalamic nucleus, posterior nucleus of thalamus, ventrobasal complex, zona incerta, medial lemniscus etc.) and the subthalamic nucleus were taken into consideration based on the atlases(*31, 33*).

TH immunostaining was carried out on coronal sections containing SNc and ventral tegmental area (VTA) according to the immunoperoxidase protocol described above. The number of TH+ cells per unit area was determined. For the estimation of dopaminergic cell loss, we used a method of Rice et al 2016(*35, 36*). The loss of DAergic neurons was determined by counting TH-immunoreactive cells under bright-field illumination. Each section was photographed using a 10X objective by randomly setting a fixed point in the Z axis within the section (Zeiss, LSM, Nikon Eclipse, Nikon 50i, 600 dpi uncompressed TIFF file). One to five nonoverlapping images were taken from each section to cover the entire SN/VTA complex within the section on both treated and non-treated sides. Each image was considered a “counting frame” with the bottom and right margins of the image as the forbidden lines of the counting frame. Neurons were selected for counting only if the entire cell body and the nucleus were clearly identified in the image. Neurons touching the right and bottom borders of the frame were excluded from counting(*35*). Our experimental design prevented double-counting of neurons, since each section selected for counting was at least 40 μm apart, considering the shrinkage factor of 5 in z-axis in case of gelatin-mounted sections (1 out of every 3 consecutive sections counted) (*37*).

The “cell counter” plug-in of NIH Image J software was used for the manual labeling of dopaminergic neurons in the images. (*36*).

#### 2.4.3 Statistical analysis of histological evaluation

The average cell number per unit area of each subregion was calculated and one-way ANOVA was used for statistical analysis with the GraphPad Prism 6.1 software(*35, 36*). When a significant difference was found between groups, we performed Tukey’s honest significant difference (HSD) post hoc multiple comparison test to identify pairwise differences. For all tests, p < 0.05 was considered significant.

#### 2.4.4 Electron microscopy – visualizing ultrastructural changes in DAergic neurons

Fifty μm sections were cryo-protected in 30 % sucrose in PB overnight. The next day, sections were freeze-thawed: immersed in 30% sucrose solution at 4°C overnight, then frozen over liquid nitrogen, then thawed on room temperature three times. Subsequently, tissue slices were washed in 0.1 M PB buffer. For intracellular detection of TH, the sections were washed subsequently in 0.1 M PB for 30 min, followed by washing in tris-buffered saline (TBS) and a 15 min treatment with borohydride in TBS. After washing out the borohydride with TBS, we were blocking the sections in 1 % human serum albumin (HSA; Sigma-Aldrich) in TBS for 1 h. Then, they were incubated in a solution of primary antibodies against TH (mouse, 1:8000 in TBS). After this, the sections were incubated in a secondary antibody solution containing gold-conjugated donkey anti-mouse (donkey-anti-mouse IgG (H&L) Ultra Small, Aurion; in a concentration of 1:100) diluted in Gel-BS. After intensive washes in BSA-c, the sections were treated with 2% glutaraldehyde in 0.1 M PB for 15 min to fix the gold particles into the tissue. To enlarge immunogold particles, this was followed by incubation in silver enhancement solution (SE-EM; Aurion) for 1h at room temperature. The sections were treated with 0.5% osmium tetroxide in 0.1 M PB on ice and they were dehydrated in ascending alcohol series and in acetonitrile and embedded in Durcupan (ACM; Fluka). During dehydration, the sections were treated with 1% uranyl acetate in 70% ethanol for 20 min. After this, 60-nm-thick serial sections were prepared using an ultramicrotome (Leica EM UC6) and picked up on single-slot copper grids. The sections were examined using a Hitachi H-7100 electron microscope and a Veleta CCD camera.

## 3. Results

### 3.1 Behavioral tests

#### 3.1.1 Locomotion in open field test

Fifty-four adult male C57BL6/J mice received unilateral injections of 6-OHDA in the dorsal striatum at three different doses. Overall horizontal locomotion was tested in an open field (OF) arena 1 week and 2 weeks after injections to assess early impairments potentially relevant to PD onset(*29*) (Fig.2). We measured the average speed, and the number of times mice crossed the lines of a grid across the arena (‘number of line-crossings’) during the full 5 minutes duration of the OF test(*12*).

**Figure 2.**
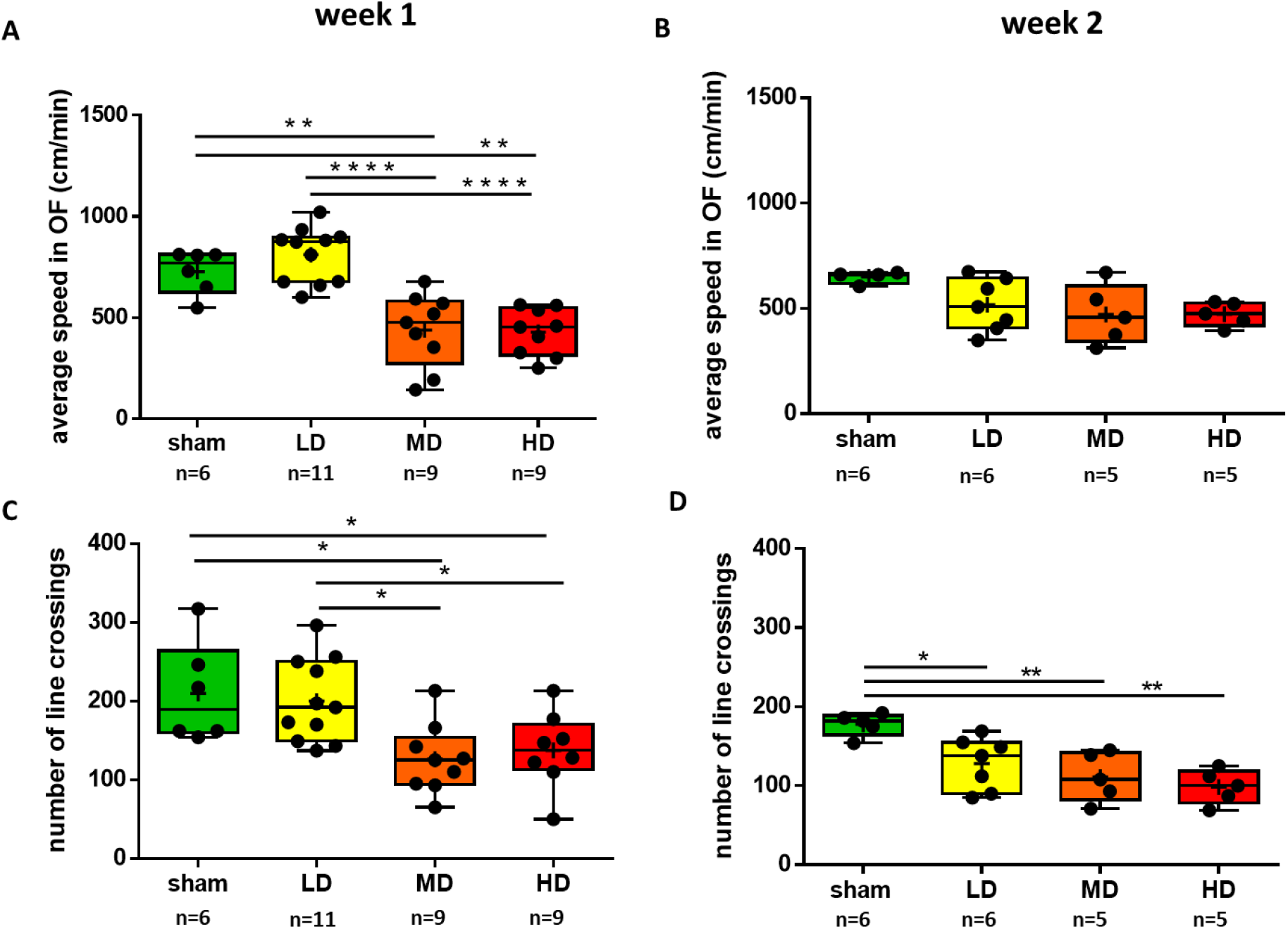
Overall horizontal locomotion showed early impairment after partial unilateral striatal 6-OHDA lesion. A, Average speed in an open field arena was measured one week after the lesion protocol. B, In half of the animals, the measurements were repeated two weeks after the lesion. C, The ‘number of line crossings’ in the OF (see Figure 1) was measured one week post lesion. D, In half of the animals, these measurements were repeated two weeks after the lesion. LD, low dose; MD, medium dose; HD, high dose. *, p < 0.05; **, p < 0.005; ***, p < 0.0005; ****, p < 0.00005.

One week after injection, we observed an acute, dose-dependent disruption of overall motility. Importantly, low dose (LD) injections did not lead to significant or even tendentious changes in average speed and number of line-crossings tested one-week post-injection (average speed, p = 0.6428; number of line crossings, p = 0.9830; one-way ANOVA, Tukey’s multiple comparation test; Fig.2A, C). At the same time, the average speed was significantly lower, and the number of line-crossings also decreased significantly in mice receiving medium dose (MD) and high dose (HD) injections compared to sham-lesioned mice (average speed: sham vs. MD, p = 0.0026; sham vs. HD, p = 0.0018; number of line crossing: sham vs. MD, p = 0.0225; sham vs. HD, p = 0.0463; one-way ANOVA, Tukey’s multiple comparation test). In accordance, we found significant differences between LD and MD, as well as LD and HD groups of mice both in mean speed and number of line-crossings tested one week after 6-OHDA injections (average speed: LD vs. MD, p < 0.0001; LD vs. HD, p < 0.0001; number of line crossing: LD vs. MD, p = 0.0170; LD vs. HD, p = 0.0333; one-way ANOVA, Tukey’s multiple comparation test). We did not find significant differences between MD and HD animals (average speed, p = 0.9988; number of line crossing, p = 0.9702; one-way ANOVA, Tukey’s multiple comparation test).

Some mice were subjected to histological analysis after the tests (see below), while others were tested again in the OF arena 2 weeks after 6-OHDA injections (Fig.2B, D). We observed a dissociation between the impact on speed of locomotion and distance covered as indexed by line-crossings. On one hand, we did not find a significant effect of the lesions on speed, however, a tendency of decrease was observed at all three doses. On the other hand, number of line-crossings showed a significant decrease in all three dose groups compared to sham-lesioned mice (sham vs. LD, p = 0.0269; sham vs. MD, p = 0.0053; sham vs. HD = 0.0011).

No rotational behavior was observed in any of the tested mice, neither one nor two weeks following drug injection

#### 3.1.2 Explorative locomotion during the first minute of the open field test

We observed that mice started to habituate to the initially new environment and move less after one minute of exploration. Therefore, to have a better insight into explorative locomotion, we performed a separate analysis of the number of line-crossings and the traveled distance during the first minute of the OF recordings (Fig.3A-B).

**Figure 3.**
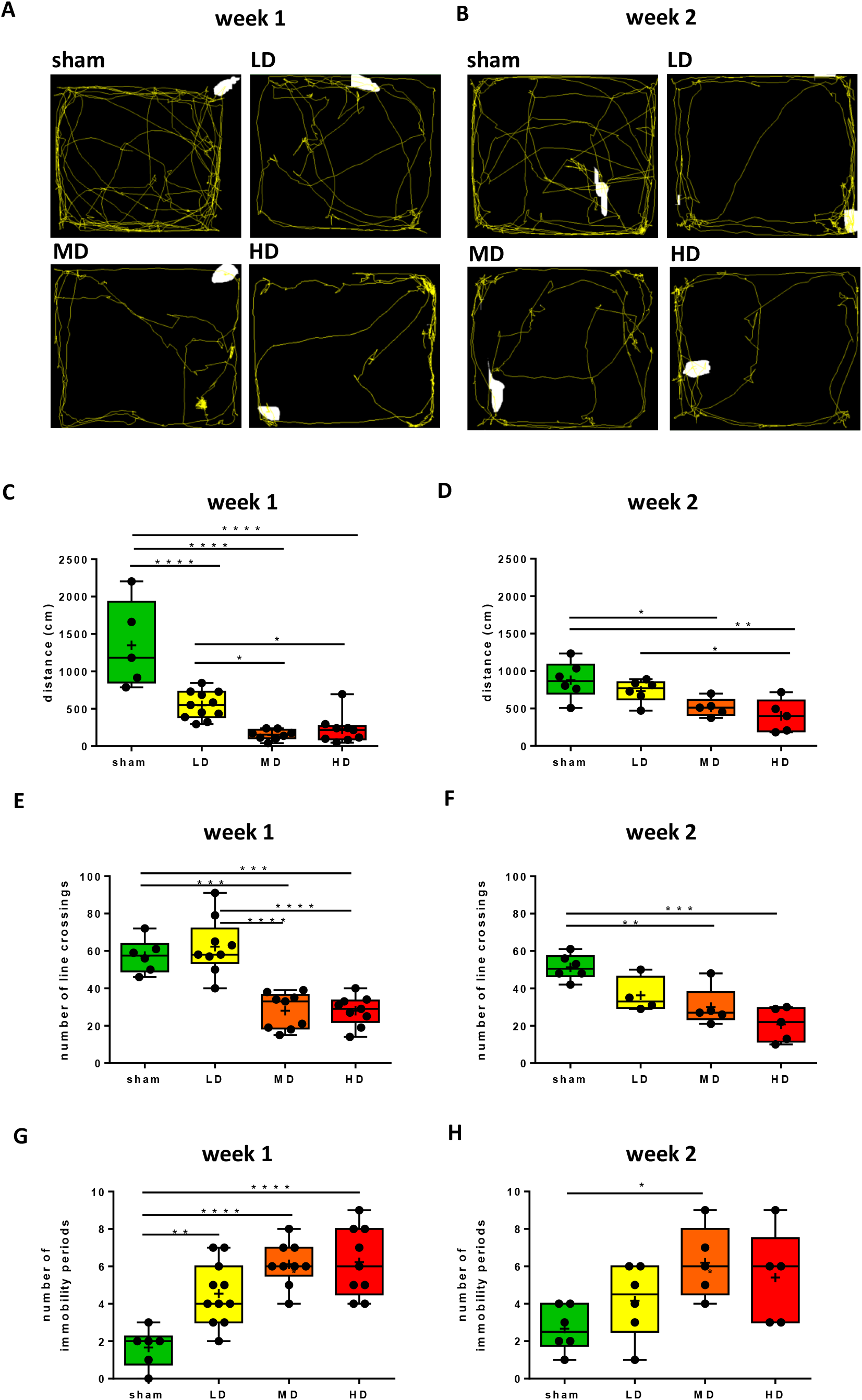
Explorative locomotion showed early impairment after partial unilateral striatal 6-OHDA lesion. A-B, Representative examples of movement trajectories during the first minute of exploration in the open field test, one (A) and two (B) weeks after 6-OHDA injection. C-D, Average distance traveled (in centimeters) during the first minute of open field exploration one (C) and two (D) weeks following 6-OHDA administration. E-F, Number of line crossings during the first minute of open field test one week (E) and two weeks (F) following 6-OHDA administration. G-H, Number of immobility periods during the first minute of open field test one week (G) and two weeks (H) following 6-OHDA administration. *, p < 0.05; **, p < 0.005; ***, p < 0.0005; ****, p < 0.00005.

We found significant differences in travelled distance when comparing sham lesioned and 6-OHDA-treated groups one week after the injections (week 1, sham vs. LD, p < 0.0001; sham vs. MD, p < 0.0001; sham vs. HD, p < 0.0001; one-way ANOVA, Tukey’s multiple comparation test). Additionally, there was a significant difference between LD and MD, as well as and LD and HD groups one week following 6-OHDA injection (LD vs. MD, p = 0160; LD vs. HD, p = 0.0330; one-way ANOVA, Tukey’s multiple comparation test; Fig.3C). We found similar, albeit less pronounced effects two weeks after the injections, when the MD and HD groups significantly differed from sham-lesioned controls, and LD vs. HD groups showed a significant difference as well (week 2, sham vs. MD, p = 0.0285; sham vs. HD, p = 0.0036; LD vs HD, p = 0.0474; one-way ANOVA, Tukey’s multiple comparation test; Fig.3D).

When we analyzed the number of line-crossings, we found that mice in the MD and HD groups crossed significantly less mesh lines than sham lesioned animals, both one and two weeks following 6-OHDA injections (week 1, sham vs. MD, p = 0.001; sham vs. HD, p = 0.001; week2, sham vs. MD, p = 0.0055; sham vs. HD, p = 0.0002; one-way ANOVA, Tukey’s multiple comparation test; Fig.3E-F). Mice in the LD group crossed significantly more lines than those in MD and HD groups one week post injection (LD vs. MD, p < 0.0001; LD vs. HD, p < 0.0001; one-way ANOVA, Tukey’s multiple comparation test). Additionally, locomotion of mice receiving low dose of 6-OHDA decreased significantly from the first week to the second week after lesion (number of line crossings: LD week 1 vs. LD week 2, p = 0.0084; Mann-Whitney U-test). As expected, no changes were seen in sham-lesioned mice from one to two weeks post-surgery (number of line crossings, week 1 vs. week 2, p = 0.6840; average speed, p = 0.0667).

#### 3.1.3 Pauses in locomotion

We noticed that mice treated with unilateral intrastriatal 6-OHDA injections showed an increase in the number of stops during locomotion. Indeed, analyzing movement trajectories in the first one minute of the OF test revealed differences between sham lesioned and 6-OHDA treated animals. Intercepting locomotion, mice stopped for grooming, rearing, or sitting still. We defined those periods which animals spent without locomotion as ‘pauses’ or ‘stops’ if the central point of mice, defined by the Animal tracker analysis software, was localized in one cm^2^ in the x-y dimension for more than two seconds. The number of pauses in all 6-OHDA treated groups showed significant differences compared to sham lesioned mice one week post injection (sham vs. LD, p = 0.0037; sham vs. MD, p < 0.0001; sham vs. HD, p < 0.0001; Fig.3G). Qualitatively similar results were observed in the smaller cohorts of mice that were re-tested two weeks after the injections (Fig.3H). While the significance of this finding is not entirely clear, the increased number of stops in 6-OHDA-injected mice may be related to bradykinesia, which is often an early symptom of Parkinson’s disease and typically leads to an increase in periods of immobility in PD patients.

#### 3.1.4 Motor coordination in the rotarod test

The rotarod test of motor coordination has been used as a sensitive indicator of dopaminergic cell loss in the SNc. We tested motor skills by measuring two parameters simultaneously, the latency to fall and the revolutions per minute (RPM) values at the time of fall from the rotating rod one week and two weeks after unilateral intrastriatal 6-OHDA injections.

We found no significant changes in the latency to fall and in the RPM values at the time of fall, either one week or two weeks post lesion (RPM at time of fall, week 1, sham vs. LD, p = 0.8109; sham vs. MD, p = 0.9979; sham vs. HD, p = 0.7142; LD vs. MD, p = 0.8571; LD vs. HD, p = 0.9948; MD vs. HD, p = 0.7573; time spend on rotarod: week 1, sham vs. LD, p = 0.8295; sham vs. MD, p = 0.9884; sham vs. HD, p = 0.6632; LD vs. MD, p = 0.9347; LD vs. HD, p = 0.9837; MD vs. HD, p = 0.7917; RPM at time of fall; week 2; sham vs. LD, p = 0.4286; sham vs. MD, p = 0.2582; sham vs. HD, p = 0.7255; LD vs. MD, p = 0.9552; LD vs. HD, p = 0.9737; MD vs. HD, p = 0.8221; time spend on rotarod: week 2, sham vs. LD, p = 0.4080; sham vs. MD, p = 0.2549; sham vs. HD, p = 0.7020; LD vs. MD, p = 0.9621; LD vs. HD, p = 0.9747; MD vs. HD, p = 0.8379; Fig.4). Given the concurrent marked impairments in the OF test, this suggests a dissociation of overall horizontal locomotion and motor coordination tested short times after partial striatal 6-OHDA-mediated lesions in mice.

**Figure 4.**
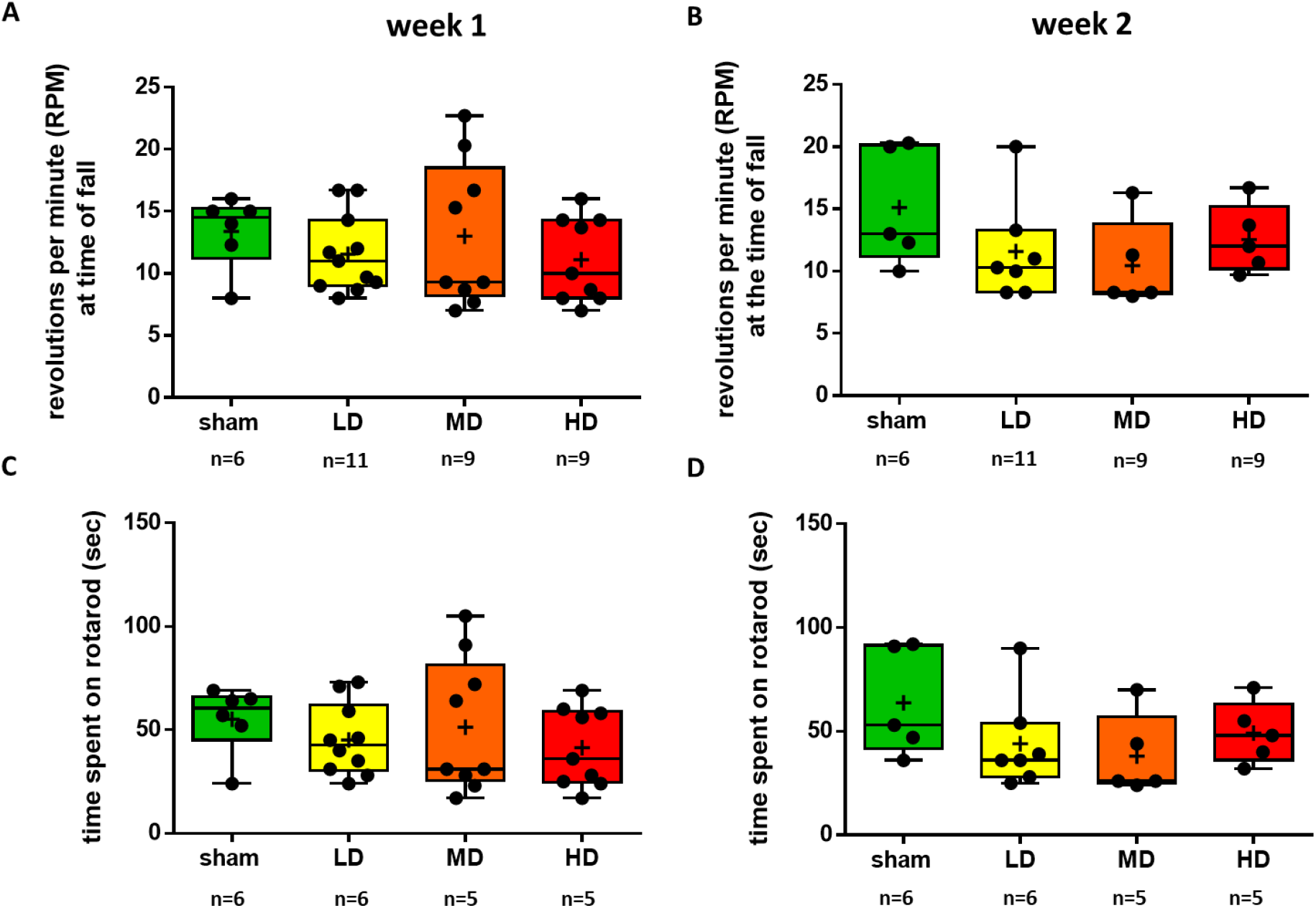
The rotarod test did not indicate and early impairment of motor coordination after partial unilateral striatal 6-OHDA lesion. A-B, The average rotation per minute (RPM) at the time of fall from the rotarod one week (A) and two weeks (B) after partial intrastriatal lesion with different doses of 6-OHDA. C-D, The latency to fall from the rotating rod one week (A) and two weeks (B) after 6-OHDA injection. LD, low dose; MD, medium dose; HD, high dose.

### 3.2 Histological evaluation

#### 3.2.1 Quantification of dopaminergic neuronal degeneration in the entire volume of the SNc one and two weeks after a single unilateral intra-striatal injection of 6-OHDA

To assess the extent of tissue damage induced by our lesion protocol, we carried out a detailed histological evaluation of retrograde degeneration of DAergic neurons in the entire volume of the SNc one and two weeks after injecting 6-OHDA in the striatum unilaterally (Fig.5). Injected hemispheres were compared with the corresponding contralateral control sides.

**Figure 5.**
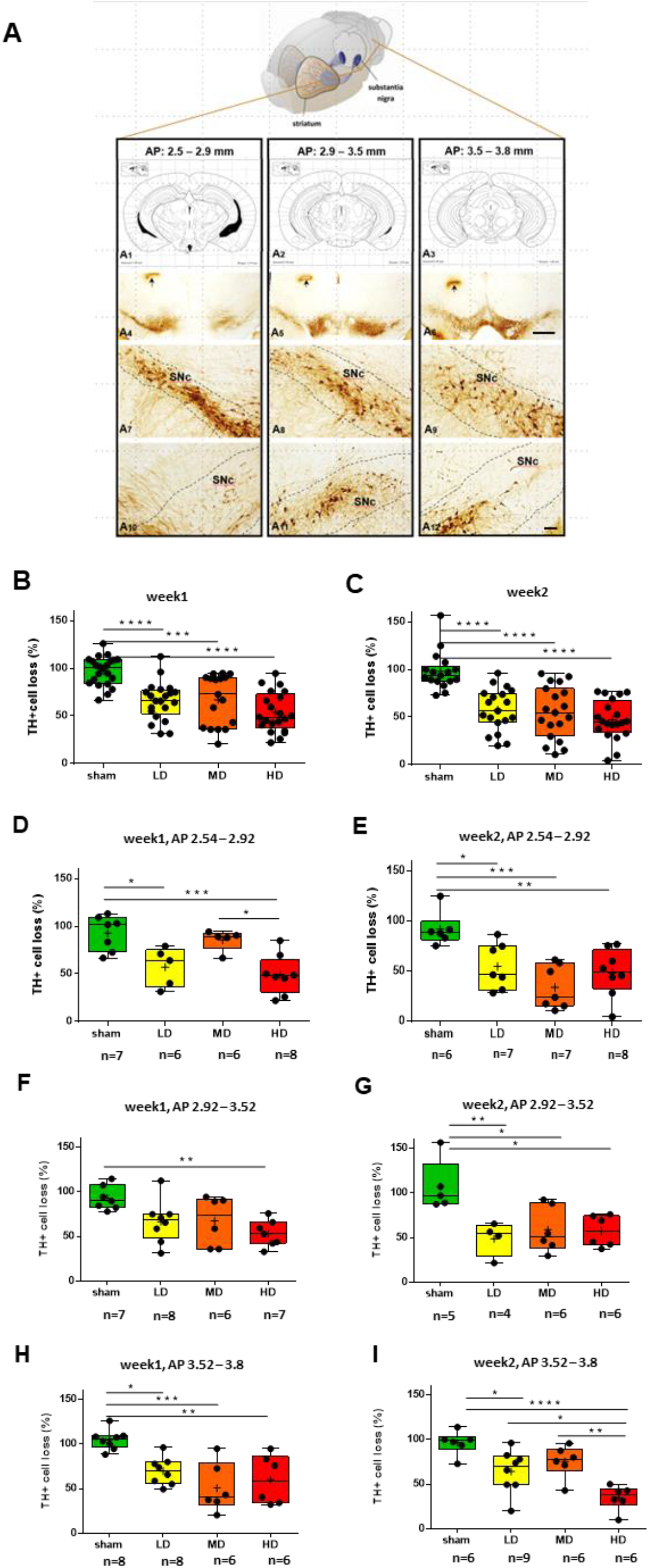
Quantification of the dopaminergic cell loss in the substantia nigra after partial unilateral striatal 6-OHDA lesion. A, The three columns represent the chosen anteroposterior levels of the mouse brain in which TH+ neurons were quantified. A_1_-A_3_, Anteroposterior ranges were selected on the basis of the Paxinos and Watson and Paxinos and Franklin atlases (range 1, AP 2.54 – 2.92 mm; range 2, 2.92 – 3.52 mm; range 3, 3.52 – 3.80 mm relative to Bregma). A_4_-A_6_, Bright field images showing DAB labelling of TH+ DAergic neurons of the substantia nigra at the three selected AP levels. Black arrows indicate tissue marks made after perfusion to label the lesioned hemisphere. A_7_-A_9_, Intact TH labelling in the substantia nigra of the control hemisphere. A_10_-A_12_, Loss of TH+ DAergic neurons on the lesioned side 2 weeks following high dose 6-OHDA injection. B-C, The percentage of TH+ DAergic neurons in the substantia nigra relative to the contralateral side one week (E) and two weeks (F) following unilateral intra-striatal 6-OHDA injection. N = 2 animals per group. Each dot represents a distinct field of view. D-E, The percentage of TH+ DAergic neurons in the substantia nigra relative to the contralateral side one week (D) and two weeks (E) following unilateral intra-striatal 6-OHDA injection quantified in the 2.54 – 2.92 mm AP range. F-G, The percentage of TH+ DAergic neurons in the substantia nigra relative to the contralateral side one week (F) and two weeks (G) following unilateral intra-striatal 6-OHDA injection quantified in the 2.92 – 3.52 mm AP range. G-H, The percentage of TH+ DAergic neurons in the substantia nigra relative to the contralateral side one week (G) and two weeks (H) following unilateral intra-striatal 6-OHDA injection quantified in the 3.52 – 3.8 mm AP range. N = 2 animals per group. Scale bar for A_4_-A_6_, 500 µm; for A_7_-A_12_, 100 µm. *, p < 0.05; **, p < 0.005; ***, p < 0.0005; ****, p < 0.00005.

We found significant neurodegeneration of DAergic neurons of the SNc following 6-OHDA injection. This was apparent already after the low-dose injection, and we did not detect a significant dose effect in dopaminergic neuron counts. The extent of dopaminergic cell loss was similar when assessed one vs. two weeks post injection surgery, demonstrating a definitive early-stage damage (mean values of number of TH+ neurons in percentage compared to control side, week1, sham, 97.25; LD, 65.56; MD, 66.97; HD, 53.73; week 2, sham, 98.14, LD, 57.59; MD, 54.95; HD, 47.12; statistical comparisons, week1, sham vs. LD, p < 0.0001, sham vs. MD, p = 0.0001, sham vs. HD, p < 0.0001, LD vs. MD, p = 0.9967, LD vs. HD, p = 0.2528, MD vs. HD, p = 0.2075; week2, sham vs. LD, p < 0.0001, sham vs. MD, p < 0.0001, sham vs. HD, p < 0.0001, LD vs. MD, p = 0.9844, LD vs. HD, p = 0.4835, MD vs. HD, p = 0.7075; Fig. 5B-C).

#### 3.2.2 Quantification of dopaminergic neuronal degeneration at three different anteroposterior levels one and two weeks after a single unilateral intra-striatal injection of 6-OHDA

In order to characterize whether the different anteroposterior (AP) levels of the SNc were differentially affected by DAergic neurodegeneration, we examined whether neuronal cell loss was present at distinct AP levels of the SNc. The entire SNc was divided into three consecutive anteroposterior subregions corresponding to three ranges of AP levels (range 1, 2.5 - 2.9 mm; range 2, 2.9 - 3.5 mm; range 3, 3.5 - 3.8 mm; Fig.5). We quantified and compared the extent of DAergic neuronal loss induced by different 6-OHDA concentrations, and after different survival times at these three different anteroposterior levels. (Fig. 5D-I).

When tested one week after injections, we observed a dose-dependent effect of 6-OHDA on SNc DAergic neurons. Specifically, we detected significant loss of DAergic neurons in the HD group at all three AP levels, whereas the LD and MD groups tested significant compared to sham-lesioned mice at only one (MD) or two (LD) AP levels (week 1, AP 2.54 - 2.92: sham vs. LD, p = 0.0168; sham vs. MD, p = 0.9121; sham vs. HD, p = 0.0009; LD vs. MD, p = 0.0997; LD vs. HD, p = 0.8745; MD vs. HD, p = 0.0116; AP 2.92 - 3.52: sham vs. LD, p = 0.0925; sham vs. MD, p = 0.1460; sham vs. HD, p = 0.0075; LD vs. MD, p = 0.9999; LD vs. HD, p = 0.6048; MD vs. HD, p = 0.6170; AP 3.54 - 3.8: sham vs. LD, p = 0.0115; sham vs. MD, p = 0.0003; sham vs. HD, p = 0.0027; LD vs. MD, p = 0.3454; LD vs. HD, p = 0.8291; MD vs. HD, p = 0.8566).

We observed pronounced DAergic neuronal loss at all three AP levels, with all three injection doses when tested two weeks after 6-OHDA injections (Fig.5B-I). Mice injected with high dose showed significantly less remaining DAergic cells at the posterior-most AP level compared to both the LD and MD group, indicating dose-dependent retrograde DAergic degeneration (week 2, AP 2.54 - 2.92: sham vs. LD, p = 0.0250; sham vs. MD, p = 0.0004; sham vs. HD, p = 0.0058; LD vs. MD, p = 0.3050; LD vs. HD, p = 0.9448; MD vs. HD, p = 0.5779; AP 2.92 - 3.52: sham vs. LD, p= 0.0078; sham vs. MD, p = 0.0146; sham vs. HD, p = 0.0118; LD vs. MD, p = 0.9096; LD vs. HD, p = 0.9402; MD vs. HD, p = 0.9996; AP 3.54 - 3.8: sham vs. LD, p = 0.0206; sham vs. MD, p = 0.2426; sham vs. HD, p < 0.0001; LD vs. MD, p = 0.6835; LD vs. HD, p = 0.0348; MD vs. HD, p = 0.0050).

#### 3.2.3 Qualitative analysis of ultrastructural changes in dopaminergic neurons

The precise mechanisms underlying neuronal injury in PD are not yet fully elucidated; however, previous studies suggested that early changes in neuronal ultrastructure of midbrain dopaminergic neurons play a key role in the pathogenesis (*38–42*). Therefore, we examined the morphological features of dopaminergic neurons in tissue samples from the SNc and the striatum by electron microscopy. Dopaminergic neurons were labeled by TH immunohistochemistry using DAB as chromogen. Pre-embedding immunogold labeling combined with a second immunoperoxidase staining was used to examine the ultrastructural changes of the degenerating DAergic neurons in the SNc.

The cellular ultrastructure of TH+ neurons showed morphological alterations in 6-OHDA treated mice. Specifically, by qualitative electron microscopic analysis, vacuoles in dendrites and axons of TH-expressing DAergic neurons were found. Additionally, swollen and vacuolized dopaminergic axons were seen in the striatum, and swollen, vacuolized mitochondria were found in the somata and proximal dendrites of the gold-labeled dopaminergic neurons. Signs of neurodegeneration were present already one week following the 6-OHDA injection. Two weeks following surgery, signs of a prominent neurodegeneration were present, including mitochondria with disintegrated membranes, as well as deformed and disintegrated dopaminergic cell bodies (Fig.6).

**Figure 6.**
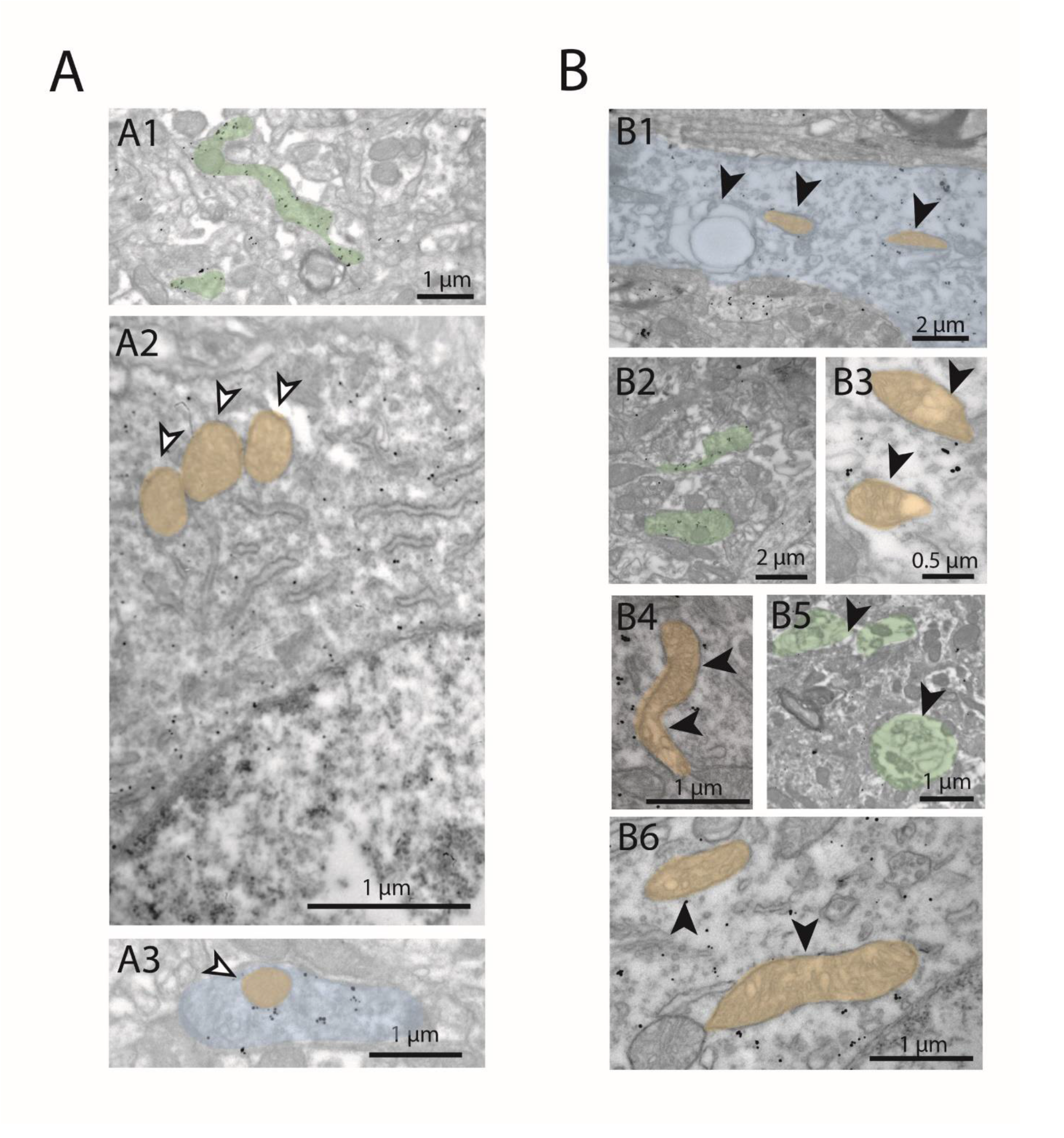
Subcellular morphological changes of dopaminergic neurons one and two weeks after striatal high-dose 6-OHDA injection. A, In sham control animals, no morphological changes were observed in the subcellular structures of SNc dopaminergic neurons in their preterminal axon segment in the striatum (A1, green), the mitochondria in their somata (A2, orange), or in their dendrites (A3, blue). Gold particles are indicating the presence of TH (immunogold staining). Note the intact inner lamellar structures of mitochondria. B, Subcellular signs of neurodegeneration were found in the injected hemisphere of 6-OHDA-injected mice. B1-B3, One week after intrastriatal 6-OHDA injection. B1, Swollen, vacuolized (black arrowhead) dopaminergic dendrite (blue) in the SNc. B2, Moderately swollen axons (green) in the striatum. B3, Vacuolized mitochondria (orange) in the soma of a dopaminergic SNc neuron. B4-B6, Two weeks after intrastriatal 6-OHDA injection. B4 and B6, Vacuolized, desintegrating mitochondria (orange) in the soma of a dopaminergic SNc neuron on the 6-OHDA-injected side. B5, Axons (green) in the striatum showing signs of neurodeneration (lamellar body formation). White arrowheads: healthy structures; black arrowheads: structures with signs of neurodegeneration.

## 4. Discussion

Early stages of PD in a graded 6-OHDA mouse model were studied in order to explore early motor symptoms and associated histological conditions at the cellular and ultrastructural level. Loss of tyrosine-hydroxylase-expressing DAergic cells and neural ultrastructural changes were characterized. Explorative behavior and motor impairments were tested by open field and rotarod behavioral assays. Our results show that mild motor impairments, characterized by a dissociation of explorative horizontal locomotion in Open Field (impaired) and motor coordination on the rotarod (intact), are detectable already at very early stages, even one week after a single low dose of striatal 6-OHDA injection.

PD leads to severe motor symptoms: tremor, rigidity, bradykinesia and postural instability(*43–46*). The key component of PD pathogenesis is a degeneration of DAergic neurons of the SNc, resulting in the deafferentation of the striatum(*7, 47*). The onset of cellular neurodegenerative mechanisms underlying PD precedes the appearance of the disease symptoms by multiple years(*14, 46, 48*): estimations date the initial presymptomatic phase 5 to 30 years prior to the clinical manifestation(*6*). Motor symptoms first appear in patients after the degeneration of 50-60% of DAergic neurons and 70-80% dopamine depletion in the striatum(*22, 44–46*).

In concert, most animal models represent an advanced phase of the disease, with substantial DAergic cell loss resulting in severe motor deficit(*24*). However, a better understanding of the early phase of PD could lead to efficient ways of early detection and intervention, potentially leading to more efficient therapeutical options. Therefore, to better understand the processes which leads to the final neuronal degeneration of the dopaminergic system in SNc, we focused on the early events of PD genesis. In this study, we examined the early changes associated with PD in a graded-dose 6-OHDA mouse model, where slight early motor impairments were correlated with mild neurodegeneration and neuronal ultrastructural changes in dopaminergic neurons. Our results show that mild ultrastructural and cellular degradation of DAergic neurons can lead to certain motor deficits in the early stage of the disease.

Toxin models achieved by intracerebral 6-OHDA injections represent a popular and powerful tool to model pathologies associated with PD. Most studies focused on rats, due to their classical role in studying the building blocks of mammalian behavior(*24, 49*). At the same time, advancements in genetic tools for mice as well as their more affordable husbandry shifted rodent research towards and increasing focus on mice. However, investigating PD models in mice has been less frequent and full-fledged 6-OHDA PD models for mice are still being developed. Injections delivered in the SNc or MFB typically result in rapid degeneration of DAergic neurons that starts as early as 24 hours within the surgery, leading to an 80% loss of DAergic neurons in 3-4 days and a complete degeneration in 3 weeks(*24, 49, 50*). To avoid a rapid and complete lesion of SNc DAergic neurons, we performed unilateral intra-striatal 6-OHDA injections at three different doses.

Dose-dependent behavioral impairments and DAergic loss of neurons and fibers were demonstrated in rats following intrastriatal injections, proposing the protocol as a suitable approach to investigate early stages of PD and to test potentially preventive interventions(*51, 52*). However, unilateral intrastriatal lesions are comparably less studied in mice. A recent line of elegant studies focusing on L-DOPA induced dyskinesia has suggested that this approach might also be viable and widely applicable for studying PD in mice(*53–57*). In accordance with these earlier studies, we demonstrated that unilateral intrastriatal graded-dose 6-OHDA injections lead to early onset dose-dependent motor deficits and DAergic lesions in mice.

The rotarod test is widely used in rodent PD models for assessing motor deficits(*24*) and a correlation between DAergic cell loss and time spent on the rotarod has been demonstrated in both rat and mouse 6-OHDA models(*58, 59*). Surprisingly, we did not find any early deficits in rotarod performance after applying our partial lesion protocol, either in RPM at the time of fall or in time latency to fall, despite the presence of DAergic degeneration. However, by carful reassessment of previous results, it appears that the strong correlation between lesion extent and rotarod performance is mostly driven by a dramatic performance drop at >80% cell loss, whereas the correlation is not obvious when considering cases with <80% cell loss, and likely not present for cases with <70% cell loss (see Fig.5. in ref(*58*)). Therefore, although the rotarod test is regarded as a sensitive indicator of DAergic degeneration in the SNc, our results suggest that it may not serve as a good marker for early deficits in rodent PD.

Although the rotarod test did not indicate an early impairment in motor coordination, motor deficits were detectable as a decrease in overall horizontal locomotion in the open field arena. Rats receiving unilateral 6-OHDA injections in the MFB(*60*), SNc(*61*) or striatum(*62*) were shown to be impaired in open field locomotion. While less studied in mice, a recent study showed similar results in a unilateral intrastriatal mouse model(*59*). On one hand, our results confirmed these earlier studies. On the other hand, the early dose-dependent impairment of horizontal locomotion that was observed in the absence of a concurrent impairment in rotarod performance indicates that the OF test may be a sensitive measure of early-stage impairment in PD mouse models. This locomotion impairment was most pronounced when we focused on exploratory locomotion in the first minute of the OF test, in line with some earlier studies showing impaired exploratory behavior in rat PD models(*63*). Medium and high dose injections led to a significant decrease in number of line crossings already when tested one week post injection, while mice injected with the lowest dose applied appeared to develop similar impairments by the second week after the surgery, shedding light on the dose-dependent time course of the early locomotion impairment.

We noticed that movement was often interrupted by grooming, rearing, or simply staying still in lesioned mice, which we defined as ‘pauses’ and quantified. This increased fragmentation of behavior may be related to problems with movement initiation and/or execution of a motor plan(*64, 65*). However, such interruptions in locomotion have not been typically associated with PD, and therefore their exact significance should be determined by future studies. It is nonetheless possible, that this phenotype is an early sign of motor deficit in PD models, that may later be masked by more severe motor impairments during disease progression.

In parallel with the behavioral characterization, we also evaluated the structural degeneration of SNc DAergic neurons by histological and electron microscopy techniques. We observed a significant loss of SNc DAergic neurons already one week after drug injection, which did not show obvious further worsening when tested two weeks post surgery. We showed by electron microscopy that gold-labeled DAergic neurons developed early signs of neurodegeneration, which can probably lead to neural cell death. The functional DAergic deficit may lead to network dysfunctions, which could accelerate disease progression(*66*), resulting in earlier cell death, as well as the emergence or worsening of clinical symptoms. Intervention should target those DAergic neurons still viable, possibly by activating neuroprotective mechanisms to slow or stop disease progression(*65, 67, 68*).

Thus, our data showed that an impairment of locomotion can be observed already one week following unilateral intrastriatal 6OHDA injection, likely in consequence of the significant decrease of SNc DAergic neurons and the ultrastructural neurodegeneration already present at that time. Studies in rats found differences in several gait parameters as early as one week following striatal 6-OHDA lesion accompanied by mild TH+ neural loss(*28*). Our partial 6-OHDA mouse model provides a useful tool to examine the early phase of PD, featuring mild early deficits with progressive characteristics, described here at the ultrastructural, histopathological, and behavioral levels. We think that models that enable studying the early phases of neurodegeneration can provide important tools to test neuroprotective strategies in animal models that might pave the way towards novel therapeutic interventions in patients with PD(*69*).

## 5. Conclusion

Due to the complex etiology of PD, its pathogenesis has not yet been fully elucidated. It is widely held that PD is a progressive and deteriorating polycentric neurodegenerative disease associated with neurotransmitter systems(*47, 70*). One main pathological mechanism of PD is the gradual degeneration and loss of DAergic neurons in the substantia nigra pars compacta, resulting in the lack of neurotransmitter DA in the basal ganglia system(*71*). Neuroprotective intervention is possible at the early stage of PD when more of the DAergic neurons and fibers remain intact. In the early stage of the disease, a targeted treatment could possibly prevent or stop the degeneration of dopaminergic neurons.

6-OHDA-mediated lesion models in rodents can be used to model distinct stages of the human pathology by varying the localization and the extent of the lesion(*24, 59*). Partial lesions of the nigrostriatal dopaminergic system might be considered analogous to the early stages of human Parkinson’s disease and can be induced reliably by intrastriatal injections of 6-OHDA. Therefore, we used this model to characterize the early signs of neurodegeneration, at which stage possible interventions might be more efficient. We described the morphological and behavioral alterations in this mouse model that may aid the quest for early diagnosis and treatment.

## Abbreviations

PD: Parkinson’s disease
6-OHDA: 6-hydroxy-dopamine
DA: dopamine
DAergic: dopaminergic
SNc: substantia nigra pars compacta
TH: tyrosine-hydroxylase
TH+: tyrosine-hydroxylase positive
LD: low dose
MD: medium dose
HD: high dose

## Acknowledgments

We thank the FENS-Kavli Network of Excellence for fruitful discussions and Dr. László Acsády for helpful comments on the manuscript. This work was supported by the ‘Lendület’ Program of the Hungarian Academy of Sciences (LP2015-2/2015), NKFIH KH125294, NKFIH K135561, the European Research Council Starting Grant no. 715043 and SPIRITS 2020 of Kyoto University to BH, and by the ÚNKP-20-3 New National Excellence Program of the Ministry for Innovation and Technology to PH. We acknowledge the help of the Nikon Center of Excellence at the Institute of Experimental Medicine (IEM), Nikon Europe, Nikon Austria and Auro-Science Consulting for kindly providing microscopy support. We thank Dr. László Acsády for providing the Nikon Eclipse confocal microscope.

